# Soluble adenylyl cyclase inhibition prevents human sperm functions essential for fertilization

**DOI:** 10.1101/2021.04.27.441671

**Authors:** Melanie Balbach, Lubna Ghanem, Thomas Rossetti, Navpreet Kaur, Carla Ritagliati, Jacob Ferreira, Dario Krapf, Lis C Puga Molina, Celia Maria Santi, Jan Niklas Hansen, Dagmar Wachten, Makoto Fushimi, Peter T. Meinke, Jochen Buck, Lonny R. Levin

**Affiliations:** Department of Pharmacology, Weill Cornell Medicine, New York City, NY; Tri-Institutional Therapeutics Discovery Institute, New York City, NY; Laboratory of Cell Signal Transduction Networks, Instituto de Biología Molecular y Celular de Rosario, Rosario, Argentina; Department of OB/GYN, Washington University School of Medicine, Saint Louis, Missouri; Institute of Innate Immunity, Biophysical Imaging, Medical Faculty, University of Bonn, Bonn, Germany

## Abstract

Soluble adenylyl cyclase (sAC: ADCY10) is essential for activating dormant sperm. Studies of freshly dissected mouse sperm identified sAC as needed for initiating capacitation and activating motility. We now use an improved sAC inhibitor, TDI-10229, for a comprehensive analysis of sAC function in human sperm. Unlike dissected mouse sperm, human sperm are collected post-ejaculation, after sAC activity has already been stimulated. Even in ejaculated human sperm, TDI-10229 interrupts stimulated motility and capacitation, and it prevents acrosome reaction in capacitated sperm. At present, there are no non-hormonal, pharmacological methods for contraception. Because sAC activity is required post-ejaculation at multiple points during the sperm’s journey to fertilize the oocyte, sAC inhibitors define candidates for non-hormonal, on-demand contraceptives suitable for delivery via intravaginal devices in females.

## Introduction

Existing family planning options are severely limited. For males, surgical vasectomy and condoms are the only available options. For females, tubal ligation provides success rates greater than 99%, but the procedure is permanent. Oral contraceptives, which are also quite effective, demand female use over prolonged periods of time and, because they are hormone-based, they carry significant side effects not easily tolerated by many women. Other effective non-surgical methods, like intrauterine devices or hormonal implants, require insertion by a doctor and suffer from similar acceptability issues. Finally, user-controlled barrier methods (e.g., diaphragms or sponges) offer added protection from sexually transmitted diseases; unfortunately, these result in failure rates greater than 13%. Thus, there is a profound need for new contraceptive strategies for males and females.

Unlike hormonal strategies, which require long-term treatment, non-hormonal contraceptive strategies allow acute interruptions to fertility. Successful on-demand contraception depends on a target that is essential for fertility and amenable to pharmacological manipulation. Bicarbonate-regulated soluble adenylyl cyclase (sAC; *ADCY10*) is the predominant, if not sole, source of the ubiquitous second messenger cAMP in sperm. Upon ejaculation, morphologically mature, but functionally immature sperm come in contact with seminal fluid. Bicarbonate in semen stimulates sAC, which activates sperm motility and initiates capacitation (i.e., the process by which sperm attain fertilizing capacity in the female reproductive tract) (reviewed in ^1-3^). In two different sAC knock out (KO) strains, males are infertile; their sperm are immotile and lack the typical hallmarks of capacitation, i.e., intracellular alkalinization, increase in protein tyrosine phosphorylation, acrosome reaction, and hyperactivated motility^4,5,6^. The dependence upon sAC for male fertility was also genetically validated in humans. Two infertile male patients were found to be homozygous for a frameshift mutation in the exonic region of *ADCY10*, leading to premature termination and interruption of the catalytic domains^7^. Similar to sperm from sAC KO mice, sperm from these patients are immotile, and the motility defect could be rescued with cell-permeable cAMP analogs. Hence, sAC is essential for sperm functions in mice and humans.

Since its initial cloning^8^, we identified multiple, chemically distinct small molecules that selectively inhibit sAC^9^. Each of these inhibitors prevented sAC-dependent functions in sperm essential for fertilization^5,10^. These pharmacological studies were performed on mouse sperm extracted from the epididymis, where sperm are stored in a low bicarbonate environment which maintains them in a quiescent state. When incubated with these freshly dissected, dormant sperm, sAC inhibitors blocked the initiation of capacitation. We now use sAC inhibitors to study human sperm. Unlike dissected mouse sperm, human sperm are collected post-ejaculation, after sAC activity has already been stimulated by the increased concentration of bicarbonate in semen relative to epididymis. Until now, it has remained unclear whether sAC activity is continuously required for steps beyond the initiation of capacitation; i.e., is cAMP synthesis necessary as sperm transit the female reproductive tract. We now demonstrate that the sAC inhibitor TDI-10229^11^ not only blocks motility, capacitation and *in vitro* fertilization in epididymis-isolated mouse sperm, it also inhibits motility and interrupts capacitation in post-ejaculated human sperm, and prevents acrosome reaction in capacitated mammalian sperm.

## Results

### TDI-10229 is an improved sAC-specific inhibitor

Previous sAC inhibitors (i.e., KH7 and LRE1) were insufficiently potent or selective to investigate their suitability as potential contraceptives. We recently developed TDI-10229^11^ and directly compared its *in vitro* potency with LRE1 both on purified human sAC protein and in a cell-based assay. LRE1 inhibited human sAC with an IC_50_ of 7.8 μM, as previously described^10^, while TDI-10229 inhibited sAC with an IC_50_ of 0.2 μM, demonstrating that our medicinal chemistry efforts significantly improved sAC inhibitory potency (Fig. 1a). To assess membrane permeability and sAC inhibitory efficiency of TDI-10229 in a cellular system, we utilized 4-4 cells, which stably overexpress sAC in a HEK293 background. Cellular levels of cAMP reflect a balance between its synthesis by adenylyl cyclases and its catabolism by phosphodiesterases (PDEs). Hence, in the presence of the non-selective PDE inhibitor IBMX, cells accumulate cAMP solely dependent upon the activity of endogenous adenylyl cyclases. Due to the overexpression of sAC, in 4-4 cells the cAMP accumulation after PDE inhibition is almost exclusively due to sAC^12,13^. TDI-10229 inhibited cAMP accumulation in 4-4 cells with an IC_50_ of 0.1 μM, which is in good agreement with its IC_50_ on pure human sAC protein and further confirms the improved potency compared to LRE1 (IC_50_ = 14.1 μM) (Fig. 1b).

**Figure 1:**
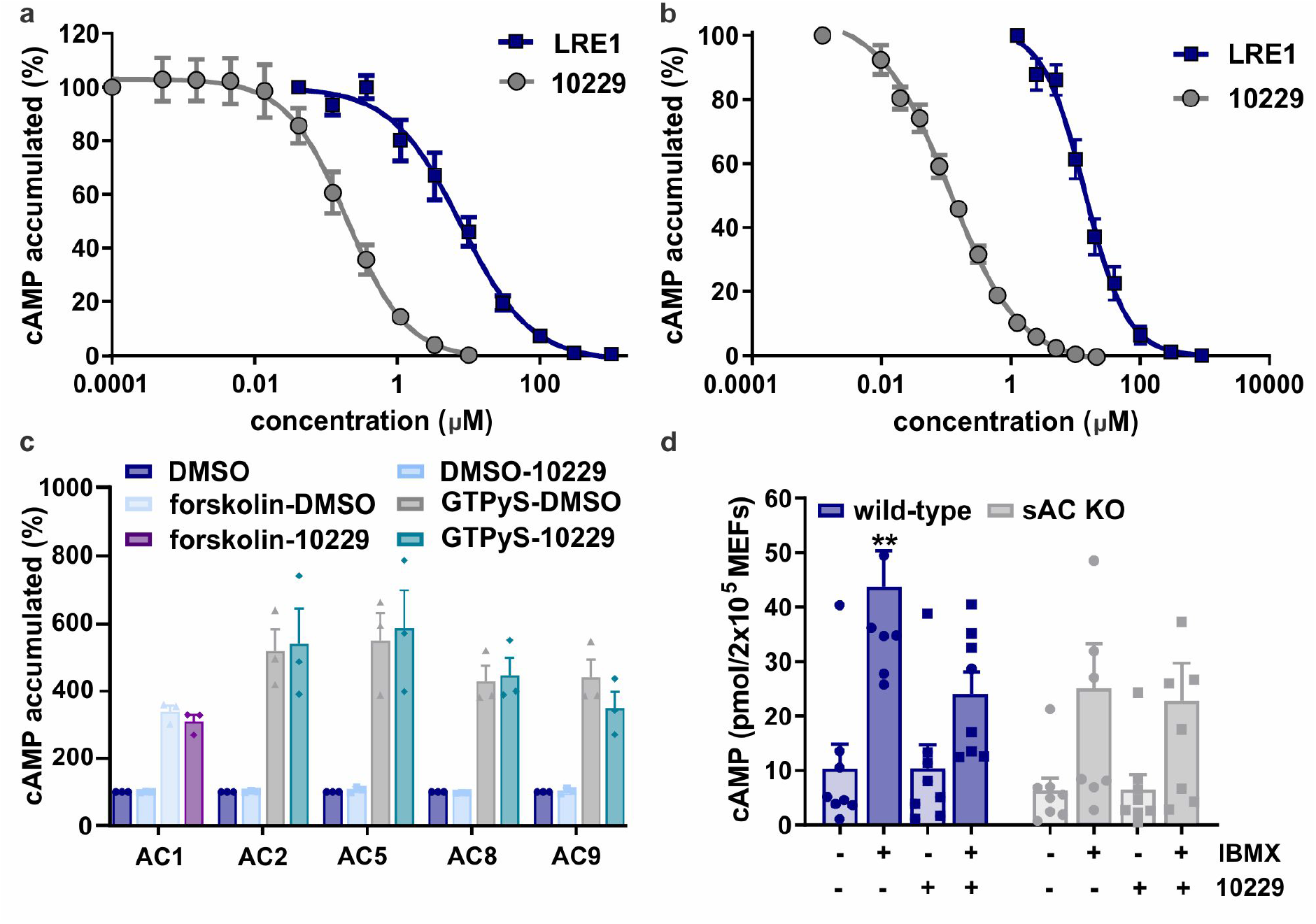
TDI-10229 is more potent than LRE1 and does not inhibit tmACs. **(a)** Concentration-response curves of LRE1 and TDI-10229 on purified recombinant human sAC protein in the presence of 1 mM ATP, 2 mM Ca^2+^, 4 mM Mg^2+^, and 40 mM HCO_3_^-^, normalized to the respective DMSO-treated control; mean ± SEM (n=5). **(b)** Concentration-response curves of LRE1 and TDI-10229 on sAC-overexpressing 4/4 cells. Cellular accumulation of cAMP measured in cells treated with 500 μM IBMX for 5 min, normalized to the respective DMSO-treated control; mean ± SEM (n=5). **(c)** AC activities of HEK 293 cell lysates overexpressing each of the indicated tmACs activated by 50 μM forskolin or 100 μM GTPyS in the absence or presence of 10 μM TDI-10229, normalized to the respective unstimulated DMSO-treated control; mean ± SEM (n=3). **(d)** Cellular accumulation of cAMP in wild-type and sAC KO MEFs treated with and without 500 μM IBMX for 10 min in the absence or presence of 5 μM TDI-10229; mean ± SEM (n=8). Differences between conditions were analyzed using (c) two-tailed, unpaired t-test or (d) one-way ANOVA compared to respective DMSO-treated control, *P<0.05, **P< 0.01, ***P<0.001, ****P<0.0001.

Mice and humans possess a single sAC gene (ADCY10) and a second, widely-expressed family of adenylyl cyclases: G protein-regulated transmembrane adenylyl cyclases (tmACs). Although tmACs are molecularly and biochemically distinct from sAC, they are the enzymes most closely related to sAC in mammalian genomes. Thus, pharmacological inhibitors should distinguish between sAC and tmACs to minimize potential off-target liabilities. To examine TDI-10229’s cross-reactivity towards tmACs, we tested whether TDI-10229 affected the *in vitro* adenylyl cyclase activities of cellular lysates each containing heterologously expressed representatives of one tmAC subclass: tmAC I, II, V, VIII, and IX. At 10 μM, 50-fold above its IC_50_ for sAC, TDI-10229 did not affect the basal nor the stimulated activities of any of the heterologously expressed tmACs (Fig. 1c). We further evaluated tmAC cross-reactivity in immortalized mouse embryonic fibroblasts (MEFs) derived from wild-type (WT) and sAC knockout (sAC KO) mice. While WT MEFs express both sAC and tmACs, the sole source of cAMP in sAC KO MEFs is the indigenous mixture of tmAC isoforms^13^. As expected, in the presence of the PDE inhibitor IBMX, WT cells accumulated more cAMP than sAC KO MEFs (Fig. 1d). And consistent with TDI-10229 selectively inhibiting sAC, TDI-10229 reduced the accumulation of cAMP in WT MEFs and was inert in sAC KO MEFs. In addition, TDI-10229 (at 20 µM, 200-fold above its IC_50_ for sAC in cells) was not cytotoxic and showed no appreciable activity against a panel of 310 kinases and 46 other well-known drug targets^11^. Thus, TDI-10229 is a potent and selective sAC inhibitor suitable for use in cellular systems.

### TDI-10229 blocks capacitation in both mouse and human sperm

Sperm are stored in a dormant state within the cauda epididymis where the bicarbonate concentration is actively maintained at ≤5 mM. Upon ejaculation, when sperm mix with seminal fluid, as well as during transit through the female reproductive tract, sperm are exposed to higher bicarbonate levels (∼25 mM)^14,15^. This bicarbonate elevation is a key initiator of capacitation; bicarbonate activates sAC which increases cAMP and protein kinase A (PKA) activity. We tested whether TDI-10229 blocks this initial signaling cascade of capacitation in mouse sperm freshly isolated from cauda epididymis. Bicarbonate-induced cAMP changes in sperm were reported to be transient^16,17^, so we first performed time course studies to establish when cAMP levels are maximal during capacitation. Bicarbonate-induced cAMP peaked at 10 minutes in mouse sperm, and TDI-10229 completely blocked the bicarbonate-dependent cAMP increases (Fig. 2a,b). As expected, cAMP levels in non-capacitating mouse sperm did not change over time, and TDI-10229 did not affect cAMP levels in sAC KO sperm (Fig. S1a).

**Figure 2:**
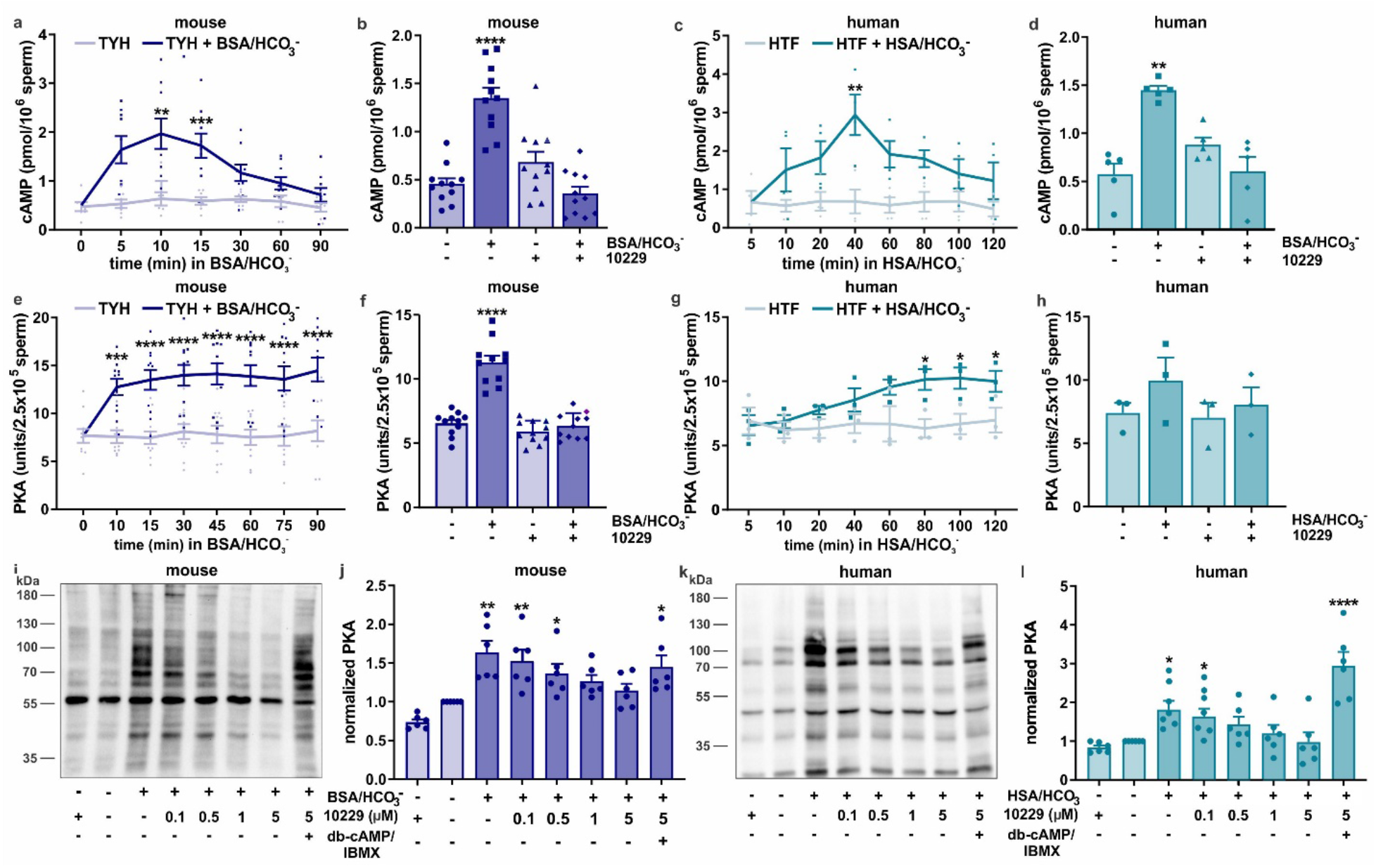
sAC inhibition by TDI-10229 prevents capacitation-induced cAMP increase and PKA activation in mouse and human sperm. **(a,c)** Intracellular cAMP levels in mouse and human sperm detected at different time points during capacitation after incubation in **(a)** TYH with 25 mM HCO_3_^-^ and 3 mg/ml BSA (0 - 90 min) or in **(c)** HTF with 25 mM HCO_3_^-^ and 3 μl/ml HSA (0 - 120 min), time-course in non-capacitated sperm is shown as control; mean ± SEM (n≥5). **(b,d)** Intracellular cAMP levels in **(b)** mouse (at 10 minutes) and **(d)** human sperm (at 40 min) in non-capacitating or capacitating media in the absence or presence of 5 μM TDI-10229; mean + SEM (n≥4). **(e,g)** Protein kinase A activity levels in **(e)** mouse and **(g)** human sperm detected at different time points during capacitation after incubation in non-capacitating or capacitating media (mouse: 0 - 90 min, human: 0 - 120 min); mean ± SEM (n≥3). **(f,h)** Protein kinase A activity levels in **(f)** mouse (at 45 minutes) and **(h)** human sperm (at 60 minutes) in non-capacitating or capacitating media in the absence or presence of 5 μM TDI-10229; mean + SEM (n≥3). **(i,k)** Phosphorylation of PKA substrates of non-capacitated and capacitated **(i)** mouse and **(k)** human sperm in the absence or presence of different concentrations of TDI-10229, rescued with 5 mM db-cAMP/500 μM IBMX, shown are representative Western blots. **(j,l)** Quantitation of PKA substrate phosphorylation Western blots of **(j)** mouse and **(l)** human sperm, normalized to DMSO-treated non-capacitated control; mean ± SEM (n≥6). Differences between conditions were analyzed using one-way ANOVA compared to the first time point (a,b), first time-point of non-capacitated control (e,f) or DMSO-treated non-capacitated control (c,d,g,h,j,l), *P<0.05, **P< 0.01, ***P<0.001, ****P<0.0001.

Unlike mouse sperm which are extracted from the epididymis in a dormant state, human sperm are isolated post-ejaculation; therefore, sAC in human sperm has already been initially stimulated via exposure to elevated bicarbonate in semen. We examined the time course of sAC activation and cAMP generation in purified and washed sperm. Bicarbonate-induced cAMP peaked at 40 minutes in human sperm, and TDI-10229 completely blocked the bicarbonate-dependent cAMP increases (Fig. 2b,d). Similar to mouse sperm, cAMP levels in human sperm incubated in non-capacitating conditions did not change over time.

Next, we measured capacitation-induced changes in PKA activity using two different assays; we directly quantified PKA enzymatic activity against an artificial substrate, and we detected endogenous PKA phosphorylated proteins. Using the direct measurement of PKA enzymatic activity, mouse and human sperm PKA activities increased during capacitation, and, in both species, the capacitation-induced increases were completely prevented by inhibiting sAC with TDI-10229 (Fig. 2e-h). Similar to the cAMP measurements, PKA activity in both mouse and human sperm incubated in non-capacitating conditions did not change over time, and TDI-10229 did not affect PKA activity in sperm isolated from sAC KO mice (Fig. S1b). We further confirmed the efficacy of TDI-10229 on bicarbonate-induced PKA activity by measuring PKA substrate phosphorylation. In both species, TDI-10229 dose-dependently blocked the capacitation-induced increase in PKA substrate phosphorylation (Fig. 2 i-l). Cell-permeable cAMP, in combination with the PDE inhibitor IBMX, rescued the block of capacitation-induced PKA activation by TDI-10229 in both mouse and human sperm, demonstrating that the inhibition by TDI-10229 can be rescued by cAMP, the product of sAC.

During mammalian sperm capacitation, the cAMP-PKA signaling cascade elicits an increase in intracellular pH (pH_i_) (reviewed in^18,19^) and a prototypical pattern of tyrosine phosphorylation (pY)^20^. We tested whether TDI-10229 blocks these molecular hallmarks of capacitation in mouse and human sperm. Sperm alkalization controls the function of multiple proteins involved in capacitation; i.e., activation of the KSper^21^ and/or CatSper ion channels^22^. During capacitation, the basal pH of 6.7 in non-capacitated mouse sperm increased approximately by 0.3 units, similar to previously reported^23,24^ (Fig. 3a). This capacitation-induced alkalization was fully blocked by inhibiting sAC with TDI-10229. In human sperm, the intracellular pH increased from 6.8 to 7.2 due to incubation in capacitating conditions, and this increase was fully blocked using TDI-10229 (Fig. 3b). Similarly, TDI-10229 blocked the capacitation-induced increase in pY in both human and mouse sperm (Fig. 3c-f). TDI-10229 inhibition of pY was concentration-dependent and rescued by cell-permeable cAMP in combination with IBMX.

**Figure 3:**
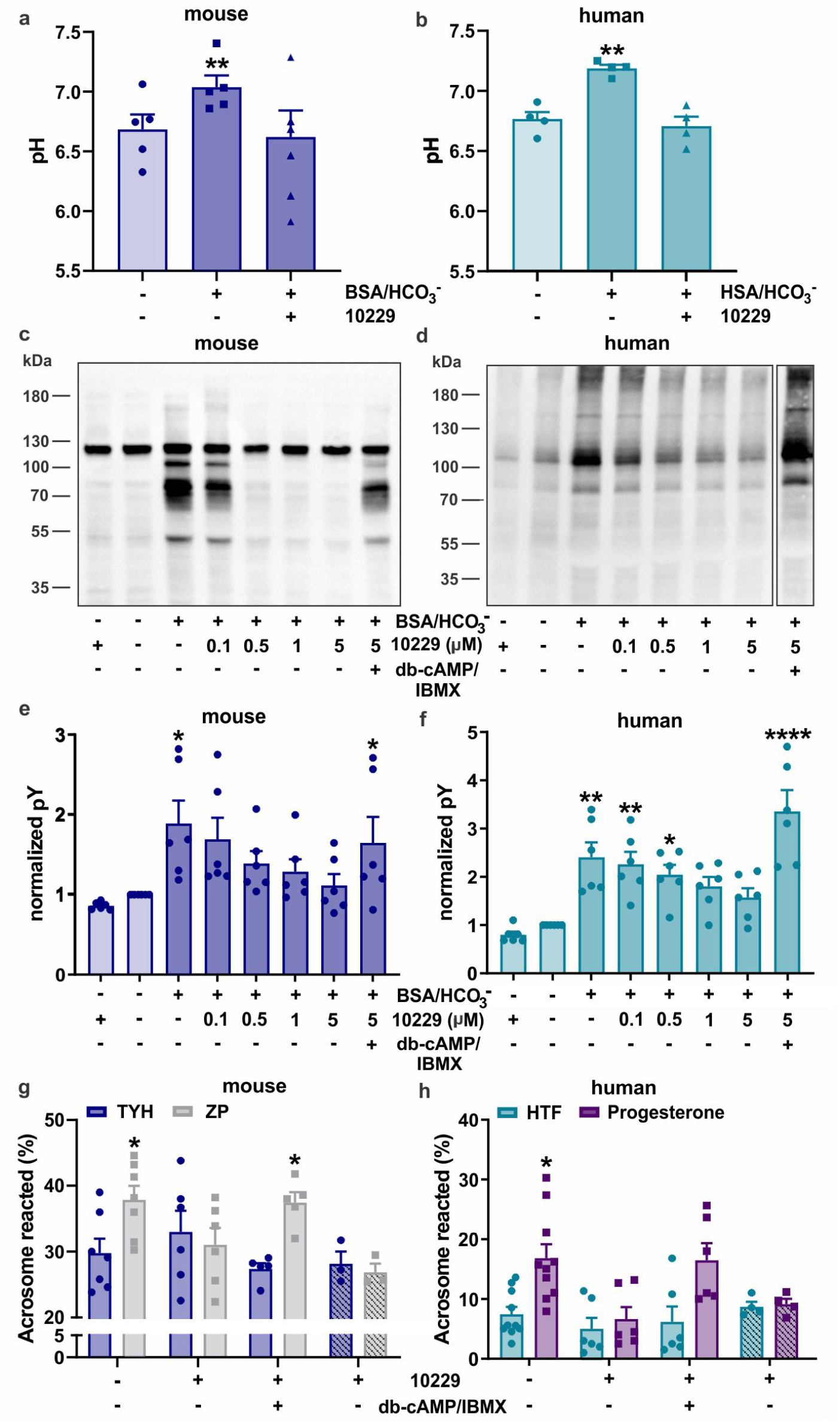
sAC inhibition by TDI-10229 prevents capacitation in mouse and human sperm. **(a-b)** Intracellular pH of non-capacitated and capacitated **(a)** mouse and **(b)** human sperm in the absence or presence of 5 μM TDI-10229; mean ± SEM (n=5). **(c,d)** Phosphorylation of tyrosine residue Western blots of non-capacitated and capacitated **(c)** mouse and **(d)** human sperm in the absence or presence of different concentrations of TDI-10229, rescued with 5 mM db-cAMP/500 μM IBMX, shown are representative Western Blots. **(e,f)** Quantitation of tyrosine phosphorylation Western blots of **(e)** mouse and **(f)** human sperm, normalized to DMSO-treated non-capacitated control; mean ± SEM (n≥6). Differences between conditions were analyzed using one-way ANOVA compared to DMSO-treated non-capacitated control, *P<0.05, **P< 0.01, ***P<0.001, ****P<0.0001.

### sAC inhibition by TDI-10229 inhibits motility of mouse and human sperm

In both mouse and human sperm, bicarbonate induced a rapid increase in flagellar beat frequency^2,25^. We characterized the flagellar beating pattern of mouse and human sperm using single sperm cells tethered via their heads to a glass surface. Analyses of flagellar beat parameters revealed that under basal conditions, both mouse and human sperm displayed a characteristic sigmoidal flagellar beating pattern, with a basal beat frequency of approximately 10 and 20 Hz, respectively (Fig. 4,5, video 1,4). Upon stimulation with bicarbonate, consistent with other mouse studies^26^, mouse and human sperm increased their overall beat frequency to greater than 35 and 25 Hz, respectively. These increases were spatially distinct between mouse and human. In mouse sperm, the increase was focused on the distal end of the tail (i.e., ≥ 60 μm from the head), while the motility of the first half of the flagellum became more restricted (i.e., ≤ 20 Hz). In contrast, in human sperm, the bicarbonate-induced increase in beat frequency was distributed over the entire flagellum. Human sperm also displayed increased curvature of their flagella.

**Figure 4:**
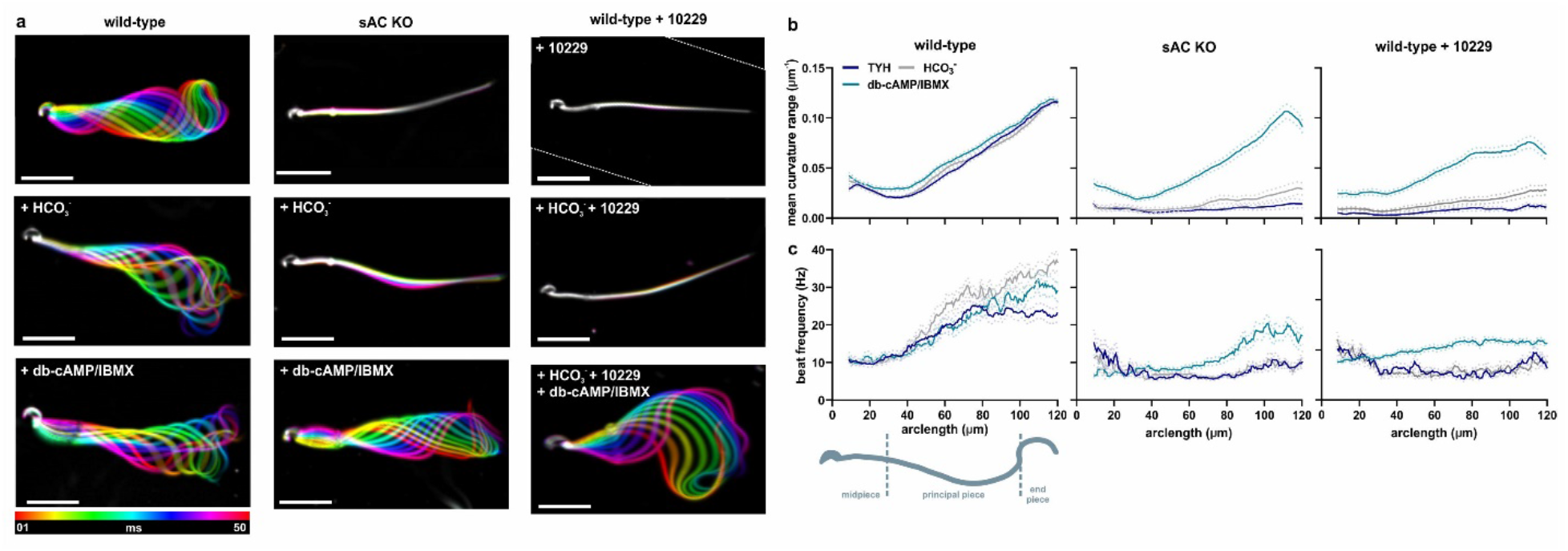
Characterization of mouse sperm beating pattern in the absence or after inhibition of sAC. **(a)** Flagellar waveform of wild-type sperm in the absence or presence of 5 μM TDI-10229 and sAC KO sperm before and after stimulation with 25 mM NaHCO_3_ or 5 mM db-cAMP/500 μM IBMX. Superimposed color-coded frames taken every 5 ms, illustrating one flagellar beat cycle; scale bar: 30 µm. **(b,c) (b)** Mean amplitude of the curvature angle and **(c)** flagellar beat frequency along the flagellum of wild-type sperm in the absence or presence of 5 μM TDI-10229 and sAC KO sperm. Solid lines indicate the time-averaged values, dotted lines the SEM, n = 3, ≥50 individual sperm from 3 different mice.

**Figure 5:**
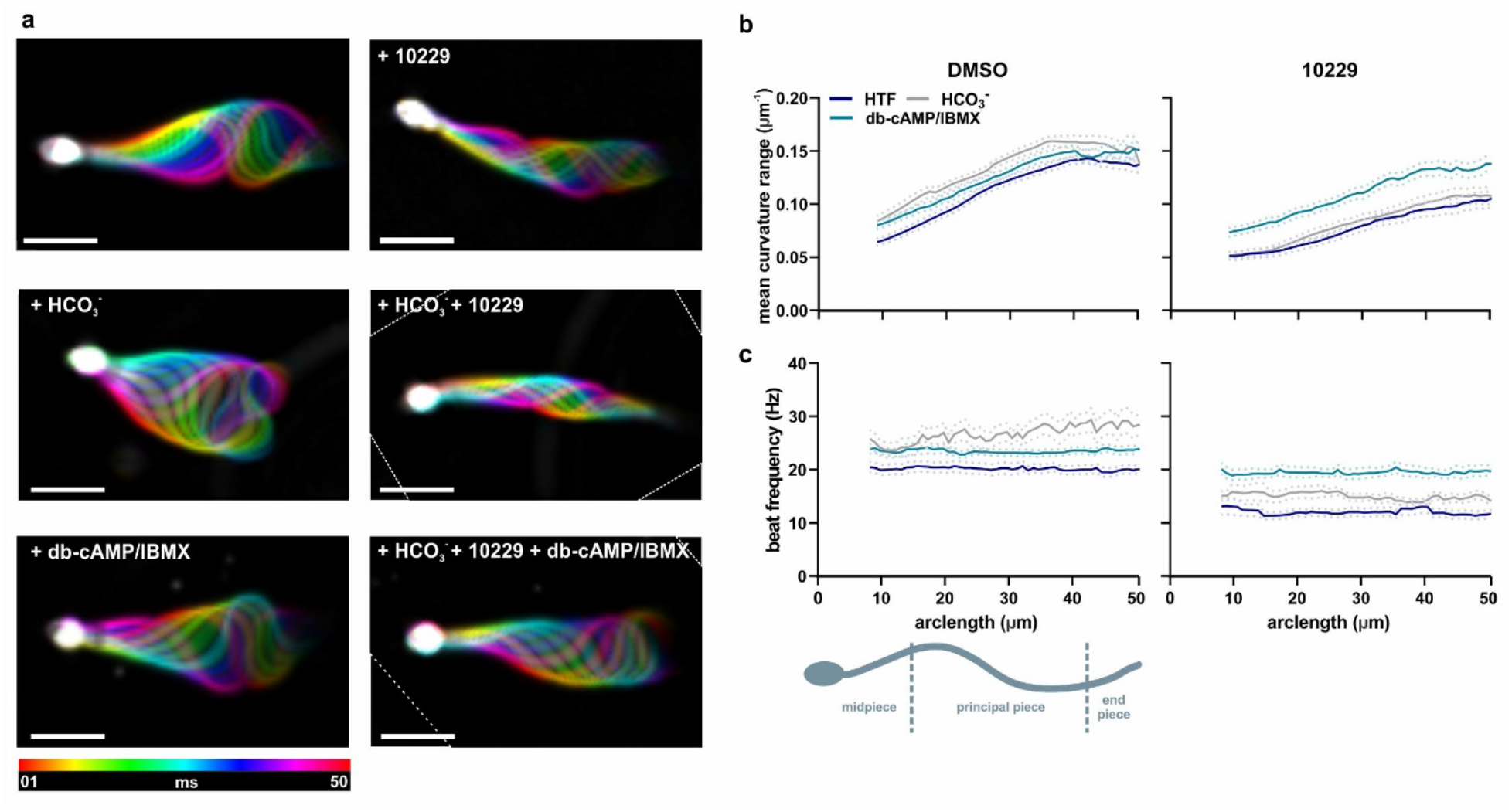
sAC inhibition by TDI-10229 prevents bicarbonate-induced changes in the flagellar beating pattern of human sperm. **(a)** Flagellar waveform of human sperm incubated in the presence of 3 μl/ml human serum albumin in the absence or presence of 0.2 μM TDI-10229 before and after stimulation with 25 mM NaHCO_3_ or 5 mM db-cAMP/500 μM IBMX. Superimposed color-coded frames taken every 5 ms, illustrating one flagellar beat cycle; scale bar: 15 µm. **(b,c) (b)** Mean flagellar beat frequency and **(c)** mean amplitude of the curvature angle along the flagellum of mouse sperm in the absence or presence of 5 μM TDI-10229 before and after stimulation with 25 mM NaHCO_3_ or 5 mM db-cAMP/500 μM IBMX. Solid lines indicate the time-averaged values, dotted lines the SEM, n = 3, ≥50 individual sperm from 3 different donors.

In sperm from sAC KO mice^4-6^ and sperm from patients predicted to have a sAC loss-of-function mutation^23^, a severe motility defect was reported. Similarly, in our hands, sAC KO sperm only showed small vibratory movements; they displayed an average beat frequency ≤ 10 Hz, and their mean curvature range along the flagellum was severely reduced (Fig. 4, video 2). When incubated with bicarbonate, the beat frequency of sAC KO sperm remained unchanged. Interestingly, the mean curvature range of sAC KO sperm ≥ 60 μm from the head slightly increased after stimulation with bicarbonate, suggesting that some aspects of mouse sperm motility might not be regulated by sAC. Incubating sAC KO sperm with cell-permeable cAMP and IBMX at least partially rescued the defects in motility, increasing both the mean curvature range and the beat frequency (Table 1).

**Table 1:** Beat frequencies of mouse and human sperm in the presence and absence of TDI-10229. Beat frequencies were averaged over the distal 10 μm of the flagellum for mouse sperm and the distal 5 μm of the flagellum for human sperm, mouse sAC KO sperm are shown as control; mean ± SEM, n≥50 individual sperm from 3 different mice/donors.

|  | Basal (Hz) | + HCO<sub>3</sub><sup>-</sup> (Hz) | + db-cAMP/IBMX (Hz) |
| --- | --- | --- | --- |
| Mouse WT | 24.3 ± 1.3 | 39.3 ± 1.8 | 29.0 ± 2.0 |
| Mouse sAC KO | 7.0 ± 0.5 | 7.7 ± 0.6 | 15.1 ± 1.5 |
| Mouse + TDI-10229 | 10.4 ± 0.8 | 11.7 ± 1.8 | 21.7 ± 0.8 |
| Human | 20.1 ± 0.8 | 27.1 ± 0.9 | 24.5 ± 0.6 |
| Human + TDI-10229 | 12.2 ± 0.8 | 14.5 ± 0.6 | 19.5 ± 0.8 |

Incubating freshly isolated WT mouse sperm with TDI-10229 reduced the mean curvature and basal beat frequency, resembling the small vibratory movements observed in sAC KO sperm (Fig. 4, video 3, Table 1). As expected for sperm in the presence of a sAC inhibitor, bicarbonate did not affect motility, and the response was largely rescued by incubation in cell-permeable cAMP and IBMX. In human sperm, TDI-10229 reduced the basal beat frequency to 15 Hz, and similar to mouse sperm, the sAC inhibitor blocked the bicarbonate-induced increase, and cell-permeable cAMP/IBMX rescued the response (Fig. 5, video 5).

### sAC inhibition by TDI-10229 blocks acrosome reaction in capacitated mouse and human sperm

The acrosome reaction is needed for successful fertilization: after the acrosome reaction, sperm can fuse their inner acrosomal membrane with the oocyte’s plasma membrane. In our final test of capacitation, as expected, blocking sperm capacitation with TDI-10229 prevented the zona pellucidae evoked increase in the percentage of acrosome-reacted mouse sperm; this response was rescued with db-cAMP/IBMX (Fig. 6a). Because zona pellucidae from human oocytes are not available, we induced acrosome reaction with the sex hormone progesterone in human sperm. As with mouse sperm, blocking human sperm capacitation with TDI-10229 prevented the progesterone-evoked increase in the percentage of acrosome-reacted sperm (Fig. 6b), and this increase was rescued with db-cAMP/IBMX. To test whether the acrosome reaction itself is dependent upon sAC, we added TDI-10229 to capacitated sperm. After incubating mouse sperm for 90 minutes and human sperm for 3 hours in capacitating media, TDI-10229 prevented the ZP- and progesterone-induced acrosome reactions, respectively.

**Figure 6:**
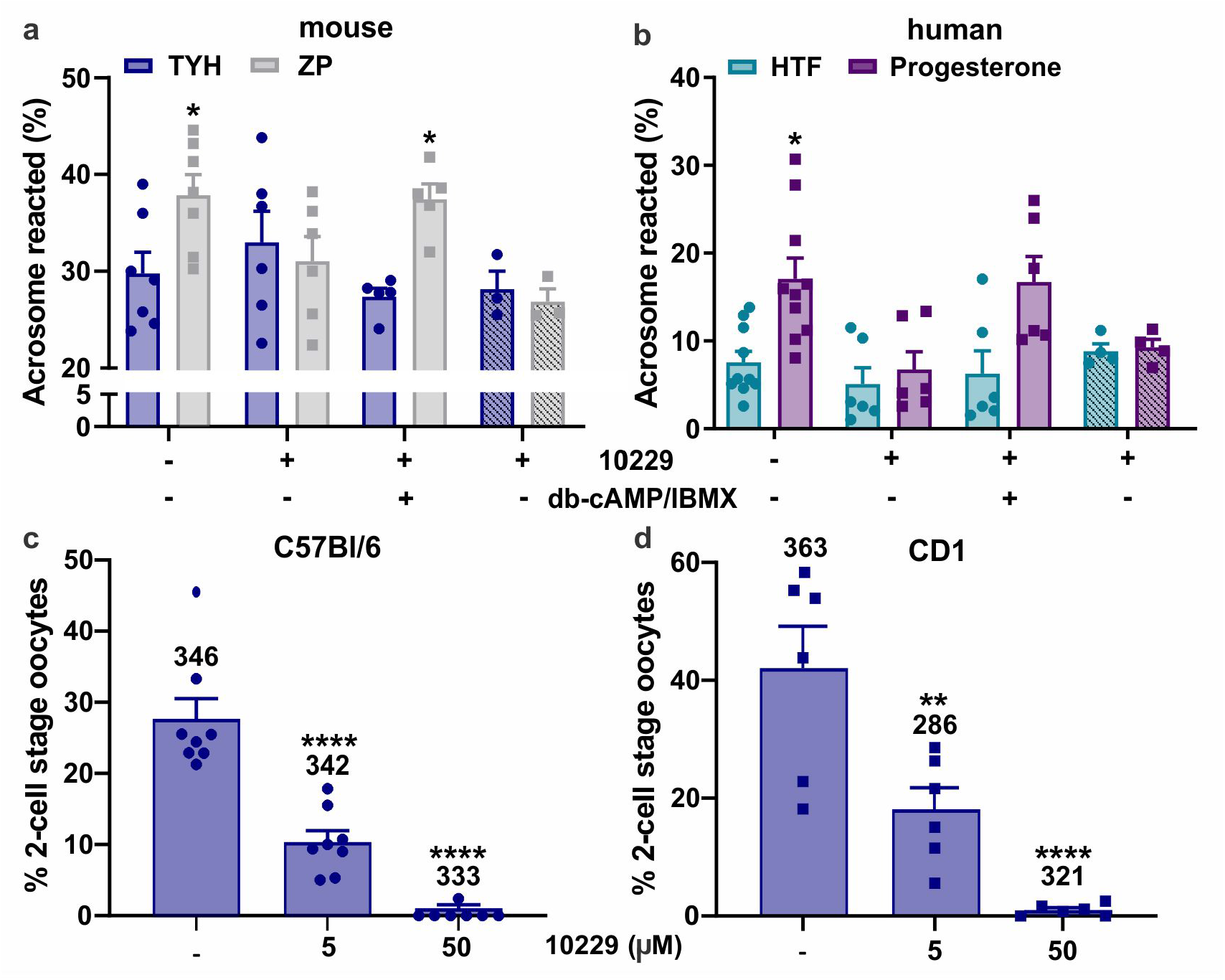
TDI-10229 blocks acrosome reaction post capacitation and fertilization of mouse sperm *in vitro*. **(a)** Acrosome reaction in wild-type mouse sperm evoked by 50 isolated zona pellucidae after incubation for 90 min in capacitating media in the absence or presence of 5 µM TDI-10229, rescued with 5 mM db-cAMP/500 μM IBMX; mean ± SEM (n≥3). **(b)** Acrosome reaction in human sperm evoked by 10 μM progesterone after incubation for 180 min in capacitating media in the absence or presence of 1 µM TDI-10229, rescued with 5 mM db-cAMP/500 μM IBMX. For the striped bars, TDI-10229 was added concomitantly with zona pellucidae or progesterone; mean ± SEM (n≥4). **(c,d)** Rate of two-cell stage oocytes after incubation of **(c)** C57Bl/6 and **(d)** CD1 mouse oocytes with capacitated C57Bl/6 and CD1 sperm, respectively in the absence or presence of 5 or 50 μM TDI-10229; mean ± SEM (n=5), numbers indicate the total number of oocytes from three independent experiments. Differences between conditions were analyzed using one-way ANOVA compared to respective DMSO-treated control, *P<0.05, **P< 0.01, ***P<0.001, ****P<0.0001.

### TDI-10229 prevents mouse in vitro fertilization

Since TDI-10229 successfully blocked capacitation and decreased beat frequency of mouse and human sperm, we tested whether TDI-10229 might be able to block fertilization of oocytes *in vitro*. Because CD1 mice are more efficient maters than C57Bl/6 mice, we performed *in vitro* fertilization experiments in both mouse strains. In a concentration-dependent manner, TDI-10229 blocked mouse *in vitro* fertilization in both C57Bl/6 and CD1. 5 µM TDI-10229, which was sufficient to block molecular hallmarks of capacitation, diminished IVF by approximately 50%, but it required higher concentrations for complete blockage (Fig. 6 c,d).

## Discussion

In this study, we validate TDI-10229 as a new pharmacological tool to study sAC-mediated biology. Due to its significantly improved potency combined with sAC selectivity, TDI-10229 allowed for a comprehensive study of the role of sAC in mouse and human sperm *in vitro*. Previous sAC inhibitors did not reproduce the motility defect of sAC KO mouse sperm^5,10^, so we utilized TDI-10229 to investigate whether this discrepancy was caused by an incomplete block of sAC activity by the less potent KH7 and LRE1 or developmental defects of sAC KO sperm. TDI-10229 fully blocked the motility of mouse sperm, resulting in small vibratory movements similar to sAC KO. These observations indicate that a) previous sAC inhibitors were indeed insufficiently potent to completely switch off sAC, and b) sAC seems to regulate mouse sperm motility at multiple levels. In addition to regulating the bicarbonate-induced increase in beat frequency, sAC-generated cAMP is also required for basal motility, at least in mouse sperm. In post-ejaculated human sperm, TDI-10229 blocked the bicarbonate-induced increase in beat frequency without affecting the basal flagellar beating pattern.

For mouse sperm, it is well established that sAC regulates the initial step of capacitation, the bicarbonate-induced increase in cAMP. Consequently, as expected, TDI-10229 also inhibited the molecular hallmarks of capacitation downstream of this intracellular increase in cAMP; i.e., PKA activation, increase in intracellular pH, and enhanced pY. The role of sAC in human sperm physiology was less explored^27,28^. In this study, we demonstrate that as in mouse sperm, human sAC regulates the bicarbonate-induced increase in cAMP, PKA activation, sperm alkalization, and increase in pY. In addition, because these studies were performed in post-ejaculated human sperm, as opposed to dormant mouse sperm isolated from the cauda epididymis, these data demonstrate that sAC does not just initiate capacitation, sAC activity is continuously required during capacitation.

The role of sAC in the acrosome reaction was less clear. While one study in human sperm showed that the sAC inhibitor KH7 blocked acrosomal exocytosis^29^, another study found that sAC KO and KH7-treated mouse sperm undergo normal acrosome reaction^5^. Because sAC is required for capacitation, and because zona pellucidae or progesterone are thought to induce acrosome reaction only in capacitated sperm, inhibiting sAC (and therefore capacitation) should also prevent the acrosome reaction. Indeed, our data demonstrate that TDI-10229 prevents the zona pellucida- and progesterone-induced acrosome reaction in mouse and human sperm, respectively. These experiments do not address the question whether sAC activity is required during the acrosome reaction. Here, we establish that adding TDI-10229 to already capacitated mouse and human sperm blocks the acrosome reaction revealing that sperm require sAC activity to undergo the acrosome reaction itself.

Because TDI-10229 blocks multiple processes sperm must complete to fertilize the oocyte, it comes as no surprise that TDI-10229 inhibits *in vitro* fertilization in mice. These results validate TDI-10229 and sAC inhibitors as potential non-hormonal contraceptives. Using intravaginal devices to deliver sAC inhibitors working topically, on ejaculated sperm in the female reproductive tract, will avoid systemic exposure and limit potential side effects in the female. To be effective, sperm-targeted, intravaginally-delivered contraceptives would have to interrupt activated sperm from completing the processes necessary to reach and fertilize the oocyte. Our use of TDI-10229 confirmed that sAC-generated cAMP is needed at multiple levels throughout the fertilization process, and that adding sAC inhibitors to post-ejaculated human sperm inhibits motility, interrupts capacitation, and prevents the acrosome reaction. Thus, delivering sAC inhibitors via intravaginal devices will provide effective contraception because: (A) By blocking sperm motility, vaginal delivery of a sAC inhibitor will prevent sperm egress from the vagina. (B) In the presence of a sAC inhibitor, any sperm that escape the vagina will fail to capacitate. And (C) a sAC inhibitor would prevent acrosome reaction in any sperm which complete the journey and survive beyond the uterus.

Intravaginal delivery devices offer additional advantages over other forms of contraceptives. Intravaginal rings and films have the capacity to simultaneously deliver multiple therapeutics which affords the unique opportunity to couple sAC inhibitor contraceptives with anti-infectives. Efforts aimed primarily at developing women-controlled products against sexual HIV-1 infection have fueled a rapid growth of intravaginal drug delivery programs, mostly involving antiretroviral drugs^31^. A number of vaginal film and intravaginal ring products are in development for HIV-1 prevention, with some candidates advancing to early-stage clinical trials^32-37^. We envision that these antiretroviral agents could be coupled with sAC inhibitors, so that one product can prevent pregnancies and sexually transmitted diseases at the same time.

## METHODS

### Reagents, cell lines, and mice

3-Isobutyl-1-methylxanthine (IBMX), bovine serum albumin (BSA), dibutyryl-cAMP (db-cAMP), BCECF-AM, hyaluronidase, lectin from Arachis hypogaea FITC-conjugated (PNA-FITC), lectin from Pisum sativum agglutinin FITC-conjugated (PSA/FITC), and mineral oil were purchased from Sigma-Aldrich, nigericin from Cayman Chemical, ionomycin from Tocris, β-mercaptoethanol from Gibco, and hormones from ProSpec. PBS buffer was purchased from Corning, EmbryoMax Modified DPBS and EmbryoMax HTF from Millipore Sigma, DMEM from Thermo Fisher Scientific and FBS from Avantor Seradigm.

4-4 cells, WT MEFs, and sAC KO MEFs were generated and functionally authenticated in our laboratory as previously described^13^ and grown in DMEM + 10% FBS. All cells were maintained at 37°C in 5% CO_2_ and were periodically checked for mycoplasma contamination.

Adult CD1-ICR (Stock #: 022) male and female mice were purchased from Charles River Laboratories and allowed to acclimatize before use. *Adcy10* KO^5^ and their corresponding wildtype were in the C57BL/6J background and bred in-house. Animal experiments were approved by Weill Cornell Medicine’s Institutional Animal Care and Use Committee (IACUC).

### *In Vitro* Cyclase Activity Assay

All i*n vitro* cyclase activity assays were performed via the “two-column” method measuring the conversion of [α-^32^P] ATP into [^32^P] cAMP, as previously described^38,39^. For the *in vitro* sAC activity assays, human sAC_t_ protein^40^ was incubated in buffer containing 50 mM Tris 7.5, 4 mM MgCl_2_, 2 mM CaCl_2_, 1mM ATP, 3 mM DTT, 40 mM NaHCO_3_ and the indicated concentration of sAC inhibitor or DMSO as control. For the *in vitro* tmAC activity assays, mammalian tmAC isozymes tmAC I (ADCY1; bovine), tmAC II (ADCY2; rat), tmAC V (ADCY5; rat), tmAC VIII (ADCY8; rat), and tmAC IX (ADCY9; mouse) were transfected and expressed in HEK293 cells using the CMV promoter. Whole-cell lysates were incubated in buffer containing 50 mM Tris 7.5, 5 mM MgCl_2_, 1mM ATP, 1 mM cAMP, 20 mM Creatine Phosphate, 100 U/ml Creatine Phosphokinase, 1 mM DTT, and 15 ng/µl DNase. When indicated, 100 µM GTPγS, 50 µM Forskolin, and/or 10μM TDI-10229 were included in the buffer. To determine tmAC-specific activities, the activities of empty vector-transfected lysates were subtracted. The activity in vector-transfected HEK293 lysates was 0.6 ± 0.03 nmol cAMP/min in the presence of 100 µM GTPγS, 0.6 ± 0.20 nmol cAMP/min in the presence of 100 µM GTPγS + 10 µM TDI-10229, 0.5 ± 0.02 nmol cAMP/min in the presence of vehicle, and 0.6 ± 0.14 nmol cAMP/min in the presence of 10 µM TDI-10229.

### Cellular cAMP accumulation assay

sAC_t_-overexpressing 4-4 cells were seeded at a concentration of 5 × 10^6^ cells/ml in 24-well plates the day before the assay in DMEM with 10 % FBS. The next day, the media was replaced with 300 µl fresh media. Cells were pretreated for 10 min with the respective inhibitor at the indicated concentrations or DMSO as control, followed by the addition of 500 µM IBMX for cAMP accumulation. After 5 min, the media was removed and the cells lysed with 250 µl 0.1 M HCl by shaking at 700 rpm for 10 min. Cell lysates were centrifuged at 2000xg for 3 min and the cAMP in the supernatant was quantified using the Direct cAMP ELISA Kit (Enzo) according to the manufacturer’s instructions.

WT and sAC KO MEFs (3 × 10^6^ cells/ml) in suspension were divided into 300 µl aliquots and incubated at 37°C for one hour. Cells were preincubated for 10 min with 5 µM TDI-10229 or DMSO as control, followed by the addition of 150 µM IBMX for cAMP accumulation. After 5 min, 150 µl of cells were transferred to a fresh tube containing 150 µl 0.1 M HCl and lysed for 10 min. Cell lysates were centrifuged at 2000xg for 3 min and the cAMP in the supernatant quantified using the Direct cAMP ELISA Kit (Enzo) according to the manufacturer’s instructions.

### Sperm preparation

Mouse sperm were isolated by incision of the cauda epididymis followed by a swim-out in 500 µl TYH medium (in mM: 135 NaCl, 4.7 KCl, 1.7 CaCl_2_, 1.2 KH_2_PO_4_, 1.2 MgSO_4_, 5.6 glucose, 0.56 pyruvate, 10 HEPES, pH 7.4 adjusted at 37°C with NaOH), prewarmed at 37°C. After 15 min swim-out at 37°C, sperm from two caudae were combined, washed two times with TYH buffer by centrifugation at 700xg for 5 min, and counted using a hematocytometer. For capacitation, sperm were incubated for 90 min in TYH containing 3 mg/ml BSA and 25 mM NaHCO_3_ in a 37°C, 5% CO_2_ incubator.

Samples of human semen were obtained from healthy volunteers with their prior written consent. Only samples that met the WHO 2010 criteria for normal semen parameters (ejaculated volume ≥ 1.5 mL, sperm concentration ≥ 15 million/mL, motility ≥ 40%, progressive motility ≥32%, normal morphology ≥ 4%) were included in this study. Sperm were purified by “swim-up” procedure in human tubular fluid (HTF) (in mM: 97.8 NaCl, 4.69 KCl, 0.2 MgSO_4_, 0.37 KH_2_PO_4_, 2.04 CaCl_2_, 0.33 Na-pyruvate, 21.4 lactic acid, 2.78 glucose, 21 HEPES, pH 7.4 adjusted at 37°C with NaOH). 0.5 to 1 ml of liquefied semen was layered in a 50 ml falcon tube below 7 ml HTF. The tubes were incubated at a tilted angle of 45° at 37°C and 15 % CO_2_ for 60. Motile sperm were allowed to swim up into the HTF layer, while immotile sperm, as well as other cells or tissue debris, did remain in the ejaculate fraction. A maximum of 5 ml of the HTF layer was transferred to a fresh falcon tube and washed twice in HTF by centrifugation (700 x g, 20 min, RT). The purity and vitality of each sample was controlled via light microscopy, the cell number was determined using a hematocytometer and adjusted to a concentration of 1×10^7^ cells/ml. For capacitation, sperm were incubated in HTF with 72.8 mM NaCl containing 25 mM NaHCO_3_ and 3 mg/ml HSA (Irvine Scientific) for 90 min to 3 h.

### cAMP quantification

Aliquots of 2×10^6^ WT or sAC KO sperm were incubated for the indicated time in the presence or absence of 5 µM TDI-10229 in non-capacitating or capacitating TYH buffer; 0.1 % DMSO was used as vehicle control. Aliquots of 2×10^6^ human sperm were incubated for the indicated time in the presence or absence of 1 µM TDI-10229 in non-capacitating or capacitating TYH buffer; 0.1 % DMSO was used as vehicle control. Sperm were sedimented by centrifugation at 2,000xg for 3 min and lysed in 200 µl HCl for 10 min. Sperm lysates were centrifuged at 2,000xg for 3 min and the cAMP in the supernatant was acetylated and quantified using the Direct cAMP ELISA Kit (Enzo) according to the manufacturer’s instructions.

### Western blot analysis

Aliquots of 2×10^6^ WT or sAC KO sperm were incubated for 45 min (PKA Western blot) or 90 min (pY Western blot) in the presence or absence of indicated concentrations of TDI-10229 in non-capacitating or capacitating TYH buffer; 0.1 % DMSO was used as vehicle control. Aliquots of 2×10^6^ human sperm were incubated for 90 min in the presence or absence of indicated concentrations of TDI-10229 in non-capacitating or capacitating HTF buffer; 0.1 % DMSO was used as vehicle control. To rescue intracellular cAMP levels, sperm were additionally incubated in the presence of 5 mM db-cAMP and 500 µM IBMX. Sperm were washed with 1 ml PBS and sedimented by centrifugation at 2,000xg for 3 min. The sedimented sperm were resuspended in 15 μl 2x Laemmli sample buffer^41^, heated for 5 min at 95°C, supplemented with 8 µl β-mercaptoethanol and heated again for 5 min at 95°C. For Western blot analysis, proteins were transferred onto PVDF membranes (Thermo Scientific), probed with antibodies, and analyzed using a chemiluminescence detection system. Image lab (Bio-Rad) was used for densitometric analysis of Western blots.

### pH assay

Sperm intracellular pH was determined as previously described^42^. Sperm samples incubated for 60 min in the presence or absence of TDI-10229 in non-capacitating or capacitating TYH or HTF buffer were incubated in the dark for additional 10 min with 0.5 μM BCECF-AM (mouse sperm) and 0.1 μM BCECF-AM (human sperm). To remove excess dye, samples were washed with the respective buffer by centrifugation at 700xg for 5 min and resuspended in non-capacitating TYH or HTF with and without bicarbonate and with or without TDI-10229. For each condition, high potassium-buffered solutions were used to calibrate the pH. 5 µM nigericin was added to each condition to equilibrate the intracellular and extracellular pH and to create a pH calibration curve. Fluorescence of BCECF was recorded as individual cellular events on a FACSCanto II TM cytometer (Becton Dickinson) (Ex: 505 nm, Em: 530/30 nm). Sperm intracellular pH_i_ was calculated by linearly interpolating the median of the histogram of BCECF fluorescence of the unknown sample to the calibration curve.

### Isolation of mouse *zone pellucida*

For *zonae pellucidae* isolation, female mice were superovulated by intraperitoneal injection of 10 I.U. human chorionic gonadotropin 3 days before the experiment. 14 h before oocyte isolation, mice were injected with 10 I.U. pregnant mare’s serum gonadotropin. Mice were sacrificed by cervical dislocation and oviducts were dissected. Cumulus-enclosed oocytes were separated from the oviducts and placed into TYH buffer containing 300 μg/ml hyaluronidase. After 15 min, cumulus-free oocytes were transferred into fresh buffer and washed twice. Zonae pellucidae and oocytes were separated by shear forces generated by expulsion from 50 nm pasteur pipettes. Zona pellucidae were counted and transferred into fresh buffer.

### Acrosome reaction assay

For analysis of acrosomal exocytosis, 100 µl 1×10^6^ sperm were capacitated for 90 min in TYH buffer supplemented with 3 mg/ml BSA and 25 mM NaHCO_3_. 5 μM TDI-10229 was added with capacitating buffer; 1 % DMSO was used as vehicle control. Acrosome reaction was induced by incubating mouse sperm with 50 mouse zona pellucida and human sperm with 10 μM progesterone for 15 min at 37 °C. The sperm suspensions were sedimented by centrifugation at 2,000xg for 5 min and the sedimented sperm were resuspended in 100 µl PBS buffer. Samples were air-dried on microscope slides and fixed for 30 minutes in 100% ethanol at RT. For acrosome staining, mouse and human sperm were incubated for 30 min in the dark with 5 μg/ml PNA-FITC and 5 μg/ml PSA-FITC in PBS. Sperm were counterstained with 2 μg/ml DAPI. After curing, slides were analyzed using a Zeiss LSM 880 Laser Scanning Confocal Microscope; images were captured with two PMTs and one GaAsP detector using the ZEN Imaging software. For each condition, at least 600 cells were analyzed using ImageJ 1.52.

### Single-sperm motility analysis

Mouse and human sperm tethered to the glass surface were observed in shallow perfusion chambers with 200 µm depth. Mouse sperm were measured in the absence of BSA since BSA affected the efficiency of TDI-10229, human sperm were measured in the presence of 3 μl/ml HSA since tethered human sperm only remain motile in the presence of HSA. An inverted dark-field video microscope (IX73; Olympus) with a 10 x objective (mouse sperm) and a 20 x objective (human sperm) (UPLSAPO, NA 0.8; Olympus) was combined with a high-speed camera (ORCA Fusion; Hamamatsu). Dark-field videos were recorded with a frame rate of 200 Hz. The temperature of the heated stage was set to 37°C (stage top incubator WSKMX; TOKAI HIT). The images were preprocessed with the ImageJ plugin SpermQ Preparator (Gaussian blur with sigma 0.5 px; Subtract background method with radius 5 px) and analyzed using the ImageJ plugin SpermQ^43^. The beat frequency was determined from the highest peak in the frequency spectrum of the curvature time course, obtained by Fast Fourier Transform.

### *In vitro* fertilization

We performed IVF experiments with both C57Bl/6 and CD1 mice. Superovulation in females was induced as described above. HTF medium (EmbryoMax Human Tubal Fluid; Merck Millipore) was mixed 1:1 with mineral oil (Sigma-Aldrich) and equilibrated overnight at 37°C. On the day of preparation, sperm were capacitated for 90 min in HTF. 100 µl drops of HTF were covered with the medium/oil mixture and 10^5^ sperm were added to each drop. Cumulus-enclosed oocytes were prepared from the oviducts of superovulated females and added to the drops. After 4 hr at 37°C and 5% CO_2_, oocytes were transferred to fresh HTF. The number of 2-cell stages was evaluated after 24 hr.

### Statistical analysis

Statistical analyses were performed using GraphPad Prism 5 (Graph-Pad Software). All data are shown as the mean ± SEM. Statistical significance between two groups was determined using two-tailed, unpaired t-tests with Welch correction, and statistical significance between multiple groups using one-way ANOVA with Dunnett correction. Differences were considered to be significant if *P < 0.05, **P < 0.01, ***P < 0.001, and ****P < 0.0001.

## ACKNOWLEDGMENT

The authors wish to thank Marc Baum for helpful insight into intravaginal delivery devices; Hannes Buck, Anna Gorovyy, Clemens Steegborn, and the rest of the Levin/Buck laboratory for helpful advice on the manuscript. The authors gratefully acknowledge the support provided by the Tri-Institutional Therapeutics Discovery Institute (TDI), a 501(c)(3) organization. TDI receives financial support from Takeda Pharmaceutical Company, TDI’s parent institutes (Memorial Sloan Kettering Cancer Center, The Rockefeller University and Weill Cornell Medicine) and from a generous contribution from Mr. Lewis Sanders and other philanthropic sources.

## AUTHOR CONTRIBUTIONS

Conceptualization: MB, PTM, LRL, JB

Methodology: MB, TR, CR, LPM, JNH

Investigation: MB, LB, TR, CR, LCPM, JF, NK

Visualization: MB

Supervision: LRL, JB

Writing – original draft: MB, LRL, JB, PTM, CMS, DW

## FUNDING

This work was funded by NIH; NICHD HD100549 & HD088571 (JB & LRL); Male Contraceptive Initiative (MB); PhD fellowship from the Boehringer Ingelheim Fonds (JNH); Germany Research Foundation under Germany’s Excellence Strategy – EXC2151 – 390873048, SPP1926, SPP1726, TRR83/SFB, FOR2743, SFB1454 (DW); FONCyT Argentina PICT 2017-3217 and PICT 2019-1779 (DK); NIH-R01HD069631 (CMS).

## COMPETING FINANCIAL INTEREST

All authors declare that they have no conflicts of interest with the contents of this article.

---

**Movie 1:** Flagellar motility of WT mouse sperm before and after stimulation with 25 mM NaHCO_3_ or 5 mM db-cAMP/500 μM IBMX.

**Movie 2:** Flagellar motility of sAC KO mouse sperm before and after stimulation with 25 mM NaHCO_3_ or 5 mM db-cAMP/500 μM IBMX.

**Movie 3:** Flagellar motility of WT mouse sperm in the presence of 5 μM TDI-10229 before and after stimulation with 25 mM NaHCO_3_ or 5 mM db-cAMP/500 μM IBMX.

**Movie 4:** Flagellar motility of human sperm before and after stimulation with 25 mM NaHCO_3_ or 5 mM db-cAMP/500 μM IBMX.

**Movie 5:** Flagellar motility of human sperm in the presence of 5 μM TDI-10229 before and after stimulation with 25 mM NaHCO_3_ or 5 mM db-cAMP/500 μM IBMX.

## Supplementary Figures

**Fig. S1:**
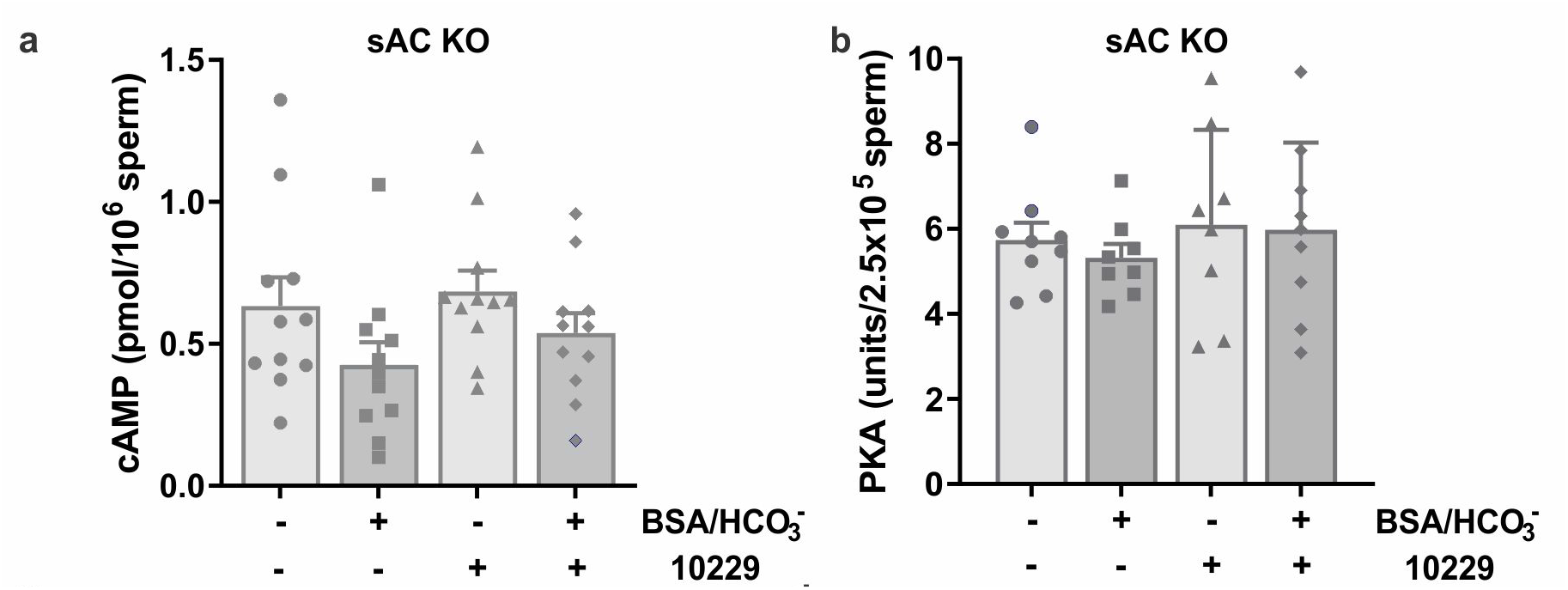
TDI-10229 is inert on sperm from sAC KO mice. **(a)** Intracellular cAMP levels in sAC KO mouse sperm after incubation for 10 min in non-capacitating or capacitating media in the absence or presence of 5 μM TDI-10229; mean + SEM (n=11). **(b)** Protein kinase A activity levels in sAC KO mouse sperm after incubation for 45 min in non-capacitating or capacitating media in the absence or presence of 5 μM TDI-10229; mean + SEM (n=9). Differences between conditions were analyzed using one-way ANOVA compared to DMSO-treated control, *P<0.05, **P< 0.01, ***P<0.001, ****P<0.0001.

## Notes

### Competing Interest Statement

The authors have declared no competing interest.

